# Enhancing Luciferase Activity and Stability through Generative Modeling of Natural Enzyme Sequences

**DOI:** 10.1101/2023.09.18.558367

**Authors:** Wen Jun Xie, Dangliang Liu, Xiaoya Wang, Aoxuan Zhang, Qijia Wei, Ashim Nandi, Suwei Dong, Arieh Warshel

## Abstract

The availability of natural protein sequences synergized with generative artificial intelligence (AI) provides new paradigms to create enzymes. Although active enzyme variants with numerous mutations have been produced using generative models, their performance often falls short compared to their wild-type counterparts. Additionally, in practical applications, choosing fewer mutations that can rival the efficacy of extensive sequence alterations is usually more advantageous. Pinpointing beneficial single mutations continues to be a formidable task. In this study, using the generative maximum entropy model to analyze *Renilla* luciferase homologs, and in conjunction with biochemistry experiments, we demonstrated that natural evolutionary information could be used to predictively improve enzyme activity and stability by engineering the active center and protein scaffold, respectively. The success rate of designed single mutants is ∼50% to improve either luciferase activity or stability. These finding highlights nature’s ingenious approach to evolving proficient enzymes, wherein diverse evolutionary pressures are preferentially applied to distinct regions of the enzyme, ultimately culminating in an overall high performance. We also reveal an evolutionary preference in *Renilla* luciferase towards emitting blue light that holds advantages in terms of water penetration compared to other light spectrum. Taken together, our approach facilitates navigation through enzyme sequence space and offers effective strategies for computer-aided rational enzyme engineering.

## Introduction

Nature has evolved enzymes as exceptional catalysts with fast turnover rates and precise specificity. Improving enzyme performance, including activity and stability, is highly desirable in various biological, medical, and industrial applications.^1^ Rational enzyme engineering provides a solution by generating variants of enzymes with improved properties, which requires a thorough understanding of enzymes and their vast variants.^2 3 4^ Computational methods have been developed to study the performance of enzyme variants,^4 5^ but accurately modeling the impact of mutations on enzyme catalytic power remains a significant challenge.^6^ Therefore, further computational tools are necessary for rational enzyme engineering.

Machine learning offers novel ways of designing enzymes.^7 8 9^ One such innovative approach involves the use of generative models to analyze evolutionarily-related protein sequences.^10 11 12 13 14^ Generative models learn a probabilistic distribution of protein sequences evolved in nature, and the probability of a specific variant correlates with its readout in deep mutation scanning experiments.^10 11^ Such correlations suggest that generative models are capable of capturing the fitness of mutations during protein evolution. Furthermore, generative models hold great promise in exploring the functional sequence space of enzymes. For instance, such models have been applied to design functional variants of different enzymes.^15 16 17 18 19^ In the majority of these enzyme engineering efforts, the tested sequences do not outperform the wild-type or natural enzymes, and even in instances where improvements were seen, a significant number of mutations have to be introduced. While these highly diverse variants with natural-like functions showcase the generative capacity of machine learning models, their practical application may face certain limitations. The introduction of too many mutations increases the cost, complexity, as well as causing poor solubility in some cases. In a clinical context, such extensively varied variants may elicit unforeseen immune responses and present ethical concerns, such as undermining genetic integrity.^20^ A critical challenge in computational enzyme engineering remains: how can we improve the enzyme to a certain performance with a limited number of mutations?

Beyond the realm of enzyme engineering, delving into natural sequences may offer insights into the intricate world of enzymes. It is well-acknowledged that both structural and functional constraints can influence evolutionary rates across protein sites.^21^ Throughout their evolutionary journey, enzymes confront numerous pressures related to their functionality and stability. Such pressures play a pivotal role in shaping enzyme evolution. For instance, mutations that enhance enzyme efficacy may be evolutionarily favored, finding their place in contemporary enzyme homologs. By harnessing generative models to extract metrics from natural sequences, we may establish links between these evolutionary metrics and the physicochemical properties of enzymes.

Through data mining of enzyme homologs with a generative maximum entropy (MaxEnt) model, we have recently illuminated the intricate evolution-catalysis relationship. Specifically, we have shed light on how enzyme sequences evolving in nature are shaped by their physicochemical characteristics, such as activity and stability, both of which are deeply intertwined with enzyme catalysis.^22^ The MaxEnt model takes into account pairwise epistasis and assigns statistical energy for each protein or mutant sequence in the multiple sequence alignment (MSA) as the evolutionary metric. For enzymes that catalyze diverse chemical reaction classes, the statistical energy corresponds with enzyme activity when the mutation is in the active center, and with stability when the mutation is situated within the scaffold. Thus, our investigations suggested key regions within enzymes that are primarily influenced by distinct evolutionary pressures.^22^ Evolution seems to prioritize the enzyme’s active center, the hub of chemical activity, and its scaffold for enhancing activity and ensuring stability, respectively. This viewpoint is far from straightforward, particularly when there are ongoing debate about whether enzymes evolved to exhibit high activity in nature.^23^ Nonetheless, the practical applicability of these theoretical insights in engineering high-performance enzymes still remains an area of exploration; the extent of the engineering success will also offer a rigorous assessment of these insights.

In this study, Renilla luciferase (RLuc) was selected as our testbed to further scrutinize the evolution-catalysis relationship, and to assess its utility in enzyme engineering. Luciferases represent a class of enzymes that catalyze the oxidation of luciferin, subsequently resulting in bioluminescent emission (Fig 1A-B).^24 25 26^ Distinctly more complex than other enzymes we have previously investigated,^22 27^ luciferases are subject to a diverse array of evolutionary pressures, encompassing factors such as catalytic efficiency, protein stability, and bioluminescence. This complexity transforms the engineering of luciferase into a multifaceted and intricate endeavor. Notably, our findings indicate that harnessing the evolution-catalysis relationship can inform rational luciferase engineering, leading to a success rate of approximately 50% in enhancing either activity or stability via the introduction of a single mutation. Beyond successful enzyme engineering, our study also revealed that RLuc has evolved to emit blue light, which has superior water penetration compared to other regions of the light spectrum. Collectively, this research underscores the potential of evolutionary data for effectively enhancing biological functions.

**Fig 1.**
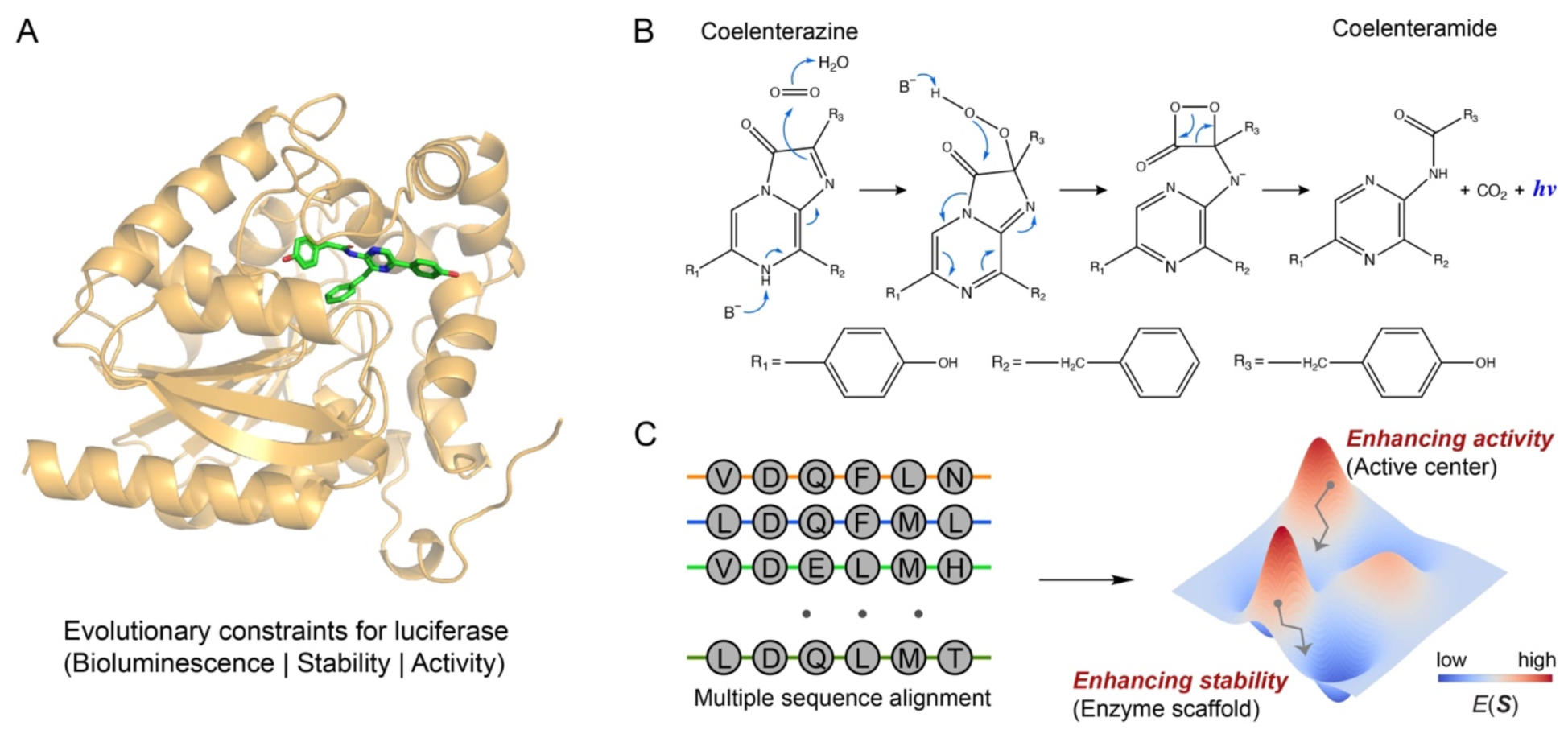
The generative maximum entropy model for luciferase. (A) Structure of RLuc with coelenterazine as luciferin. PDB ID used in rendering the structure is 6YN2 with substrate docked. (B) Scheme of the chemical reaction catalyzed by RLuc. Coelenterazine undergoes oxidation to coelenteramide with oxygen acting as the oxidant, and a photon of blue light is released as a result. (C) Rationally enhancing enzyme activity/stability guided by the generative MaxEnt model. The statistical energy *E*(***S***) as an evolutionary metric reflects evolutionary pressures. For variants with mutated residues in the active center and enzyme scaffold, *E*(***S***) highly correlates with enzyme activity and stability, respectively.^22^ Thus, enzyme performance can be improved by optimization on the evolutionary landscape.

## Results

### Generative modeling of natural luciferase sequences

Generative models analyze evolutionarily-related protein sequences and provide a probabilistic model to capture the effects of mutations in natural evolution. The probability associated with a certain sequence or variant has been connected to protein fitness.^10 11^ Among these models, the MaxEnt model, which is based on information theory, has demonstrated a strong generative capacity.^28^ The statistical energy of sequence ***S*** as derived from the model is expressed as

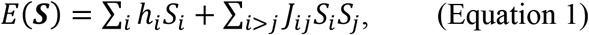

where *h*_*i*_ and *J*_*ij*_ are parameters, *S*_*i*/*j*_is amino acid at residue position *i*/*j* in the sequence. The *E*(***S***) follows the Boltzmann distribution

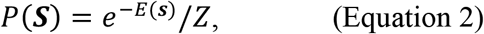

where *Z* = ∑_***S***_ *e* ^− *E*(*s*)^ is the normalization constant. The derivation and parameterization details of MaxEnt for enzymes can be found in Ref ^22^. In this context, the MaxEnt model learned from MSA describes an energy landscape as shown in the schematic picture (Fig 1C). A variant with a lower *E*(***S***) value is more likely to be found in nature, suggesting it has specific evolutionary advantages.

The complex nature of enzymes, with intertwined evolutionary pressures, means that a direct correlation between *E*(***S***) and individual parameters is not assured. This intricacy stems from the interplay of various physicochemical properties that govern enzyme function, and the multifaceted relationship between specific enzyme activities and protein thermostability.^29–31^ Such a convoluted landscape suggests that relying solely on *E*(***S***) as a predictor for various enzyme properties might be oversimplistic. However, based on our investigations on multiple enzymes, we have observed that *E*(***S***) (or *P*(***S***)) for variants with mutations in the active center and enzyme scaffold tend to correlate with enzyme activity and stability, respectively.^22^ This connection between evolution and catalysis suggested a structured approach to enhance different enzyme properties by optimizing *E*(***S***), particularly targeting distinct enzyme regions (Fig 1C).

Here we applied the MaxEnt model to luciferase for enzyme engineering. We used RLuc from *Renilla reniformis* (UniProt ID: P27652) as the target sequence and searched the UniRef90 database (release 2021_03) to construct the MSA. A length-normalized bit score threshold of 0.7 was applied. We obtained 1775 homologous sequences, which is sufficient to parameterize the model. Parameters were optimized using statistics obtained from the MSA, including the probability of a specific amino acid at a single residue and the probability of a specific pair of amino acids between two residues. Once parameterized, the statistical energy *E*(***S***) for each sequence or variant was calculated. The statistical energy was shifted so that the wild-type has a value of zero, which does not affect any analysis here. We then correlated *E*(***S***) with available biochemical data of luciferase. This correlation aimed to elucidate the association between evolution and catalysis, which could potentially shed light on nature’s strategy for generating luciferase.

### Correlating luciferase activity with natural evolutionary information

We first examined the correlation between luciferase activity and *E*(***S***), using it as an indicator of natural evolutionary information. RLuc has been extensively engineered as a crucial reporter in live-cell imaging, with 26 variants identified through consensus design.^32 33^ These variants primarily consist of double mutations, with C124A being present in all of them (Table S1). The average distance between the substrate and the mutated residues was determined for each variant, and the variants were categorized as belonging to the active center or enzyme scaffold based on an 8.5 Å cutoff.^22^ For variants classified as the active center, their activity has a significant correlation with *E*(***S***), as determined by a Pearson correlation value of -0.69 (*p*-value = 0.057) (Fig 2A). However, this correlation noticeably declines for variants with mutations on the enzyme scaffold, with a Pearson correlation of -0.25 (*p*-value = 0.29). It should be noted that the luminometers used in the measurement were calibrated to absolute units (photons/s),^32 33^ which implies that the correlation established pertains to enzyme activity and not bioluminescence.

**Fig 2.**
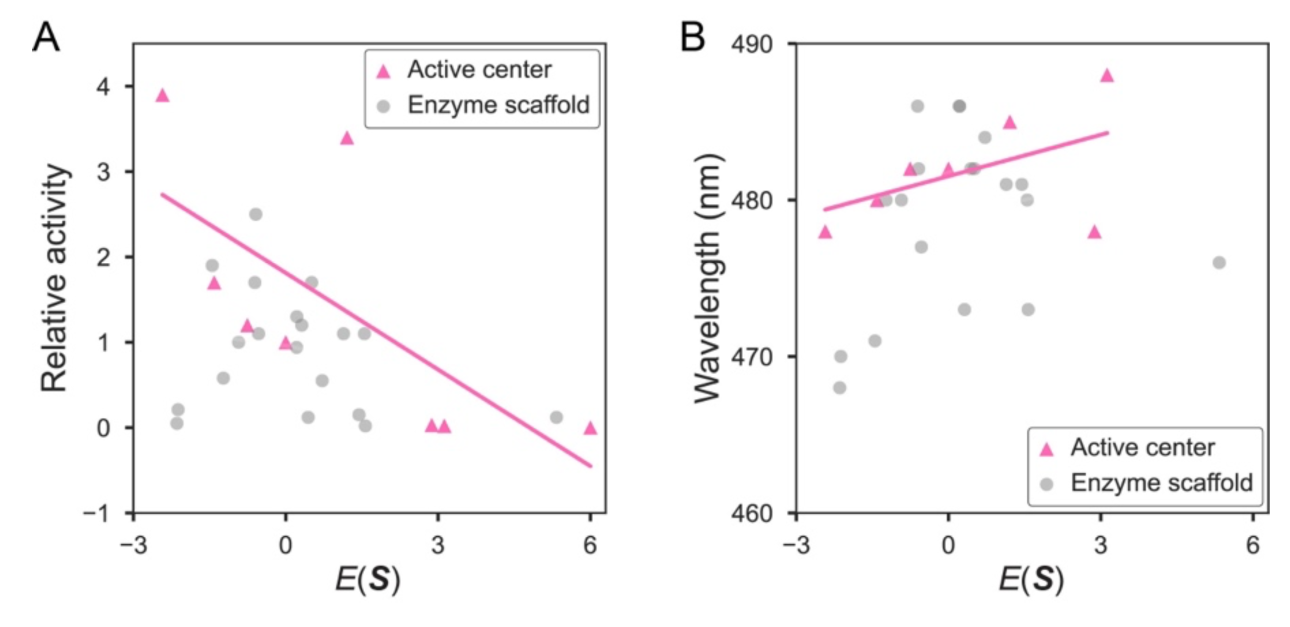
Correlations between the statistical energy *E*(*S*) and biological activity for luciferase. (A) Luciferase activity relative to wild-type. (B) Emission peak of luciferase bioluminescence.

These findings indicate that enzyme activity considerably shapes the evolution of active center compared to enzyme scaffold. The 8.5 Å cutoff value approximately distinguishes two residue shells from the substrate and places them within the active center. The first-shell residues around substrate are directly involved in substrate positioning and catalyzing reactions. There is also an increasing body of evidence to suggest that second-shell residues play a key role in regulating enzyme activity.^34^ Thus, residues within the active center have a direct impact on enzyme catalysis and are expected to be mainly shaped by biological activity during natural evolution.^22^

Intrigued by the patterns observed in catalytic efficiency, we delved deeper to discern if natural evolutionary data could also shed light on the bioluminescent attributes of luciferase. This phenomenon of bioluminescence is not just a standalone feature; it is an outcome of evolutionary pressures acting on the luciferase-mediated reactions. Parameters like emission spectrum, inactivation time, among others, play pivotal roles in shaping bioluminescence (Table S1). Our analysis revealed a notable correlation of 0.51 (*p*-value = 0.24) between *E*(***S***) and the peak emission spectrum for mutations localized in the active center, with a more modest correlation of 0.22 (*p*-value = 0.38) in the enzyme scaffold region, as seen from Fig 2. Tackling the prediction of bioluminescence spectra is notoriously tricky in computational chemistry, given it demands insight into excited states. To our knowledge, no study till date has proposed a correlation metric adept at linking bioluminescence spectra across diverse enzyme variants. This observed positive correlation hints at a natural evolutionary tendency in RLuc to favor blue light emission, possibly because of blue light’s superior water penetration capabilities compared to other wavelengths (Fig S1).^35 36^ In a related vein, nature seems to exhibit a bias towards enzymes exhibiting prolonged inactivation phases (Fig S2). Hence, *E*(***S***), in its role as an evolutionary barometer, provides a potential means to link with an array of evolutionary pressures, targeting specific protein domains.

Besides the native reaction that converts coelenterazine, we also investigated how sensitive this approach is to variations in the reacting substrate. To optimize the spectrum for live-cell imaging, the substrate has been slightly modified. The correlation between *E*(***S***) and enzyme activity is much weaker for some modified substrates (Fig S3). Thus, the natural sequences evolved for converting coelenterazine could be sensitive to the specific substrates.

Furthermore, we extended our analysis to mutations made to RLuc8, which is an engineered variant of RLuc to have eight-point mutations (Table S2). The findings underscored that the evolutionary metric, *E*(***S***), adeptly differentiates active enzymes from inactive ones, as demonstrated in Fig S4. In a more detailed assessment, we analyzed saturation mutagenesis data at the active site residue I223 of RLuc8 (Table S3). A notable correlation of -0.71 (*p*-value < 0.001) between *E*(***S***) and enzyme activity emerged, as seen in Fig S5, aligning with observations from Fig 2A. Additionally, we observed that the mutations at this residue tend to have similar effects on converting substrate analogs (Fig S6).

### Low-dimensional embedding of generated variants

In the context of enzyme design, investigating the *E*(***S***) energy landscape enables the identification of low-energy sequences that have the potential to enhance enzyme performance, as indicated from our previous analysis.^22^ Instead of sampling the energy landscape, we systematically catalog all possible single and double mutants situated in the enzyme’s active center and scaffold. By employing this straightforward strategy, we guarantee the comprehensive consideration of all lower-order mutations, which forms the central focus of the subsequent proof-of-concept biochemistry characterization. The sequences with *E*(***S***) lower than the wild-type were collected. We obtained 220 such variants with mutated residues within 8.5 Å of the substrate and 394 variants with mutated residues more than 15.0 Å from the substrate (Table S4-S5).

We used variational autoencoder (VAE)^37^ to perform the low-dimensional embedding to characterize the relation between generated variants and natural luciferase sequences (SI Methods). The VAE model was trained with natural sequences with a two-dimensional latent space. As illustrated in Fig 3A, the natural sequences are organized into different spikes in the latent space, which is also found in previous studies, potentially related to phylogeny in evolution.^38^ The variants created through redesigning the active center or enzyme scaffold are confined to a specific local region in the latent space due to the focus on single or double mutants that are similar to the wild-type (Fig 3B and 3C). These variants for the active center and enzyme scaffold occupy different regions in the latent space, close to different spikes in the latent space.

**Fig 3.**
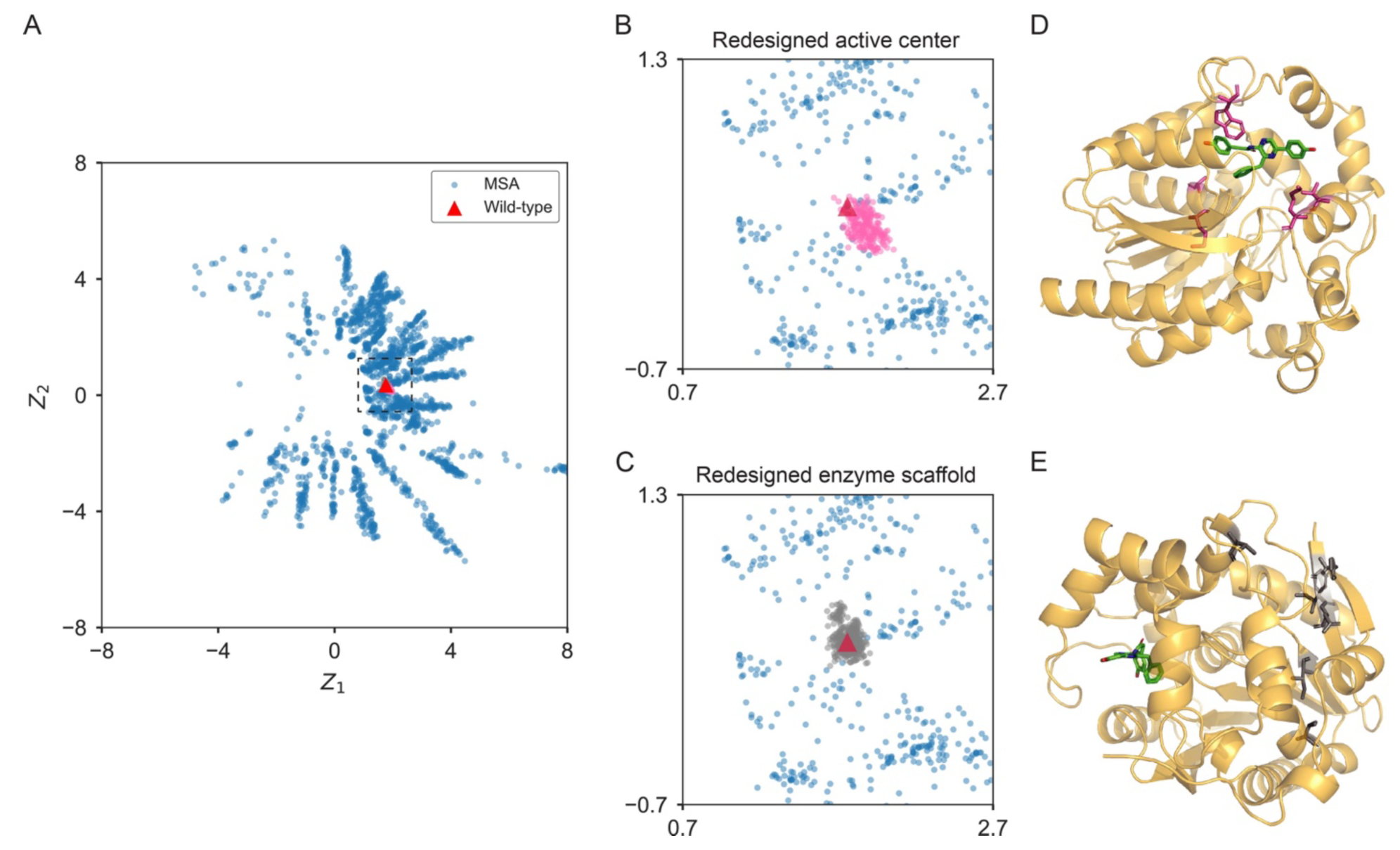
Characterization of designed luciferase variants. (A) Embedding MSA and designed sequences in a two-dimensional latent space (Z_1_, Z_2_) with VAE. (B, C) Sequences featuring a redesigned active center (B) are shown in pink, while those with a redesigned enzyme scaffold (C) are depicted in gray. (D, E) Variants that underwent testing in the experiment have their mutated residues emphasized: those in the active center (D) are colored in pink, and those on the enzyme scaffold (E) are presented in gray.

In the subsequent experiment, we selected eight single variants with the lowest *E*(***S***) values in the active center for experimental characterization. These variants are C124V, M185L, A143M, K189S, C124A, K189G, K189A, and C124G, listed in order of increasing suggestions. These mutations span four residues, all of which were successfully expressed (Fig S7). Additionally, we selected another set of eight single variants on the enzyme scaffold that also had the lowest *E*(***S***) values: I34M, I75A, E195T, F33K, V64I, N35S, Y298A, and V212S, again listed by increasing suggestions. However, Y298A and E195T mutants exhibited notably lower protein yield, and thus we characterized six mutants on the enzyme scaffold, as indicated in Fig S7. The mutated residues in the active center and on the enzyme scaffold, which were involved in the experiment, are highlighted in Figs 3D and 3E, respectively.

### Enhancing Luciferase Activity in the Active Center Validated in Experiments

For mutants in the active center, the relative activity compared to the wild-type enzyme was shown in Fig 4A. The activity of RLuc variants was determined with luminometers (as described in the SI Methods). Our results indicate that mutations in the active center have a significant impact on catalysis. Of the eight designs, half showed lowered performance compared to the wild-type activity. Notably, the A143M mutant displayed a relative activity of 1.02, and M185L, C124A, and C124G, showed notable increases in activity, with 2.01, 1.63, and 1.72 times relative activity to wild-type, respectively. Overall, about 50% of our attempts to design beneficial mutations in the active center works.

**Fig 4.**
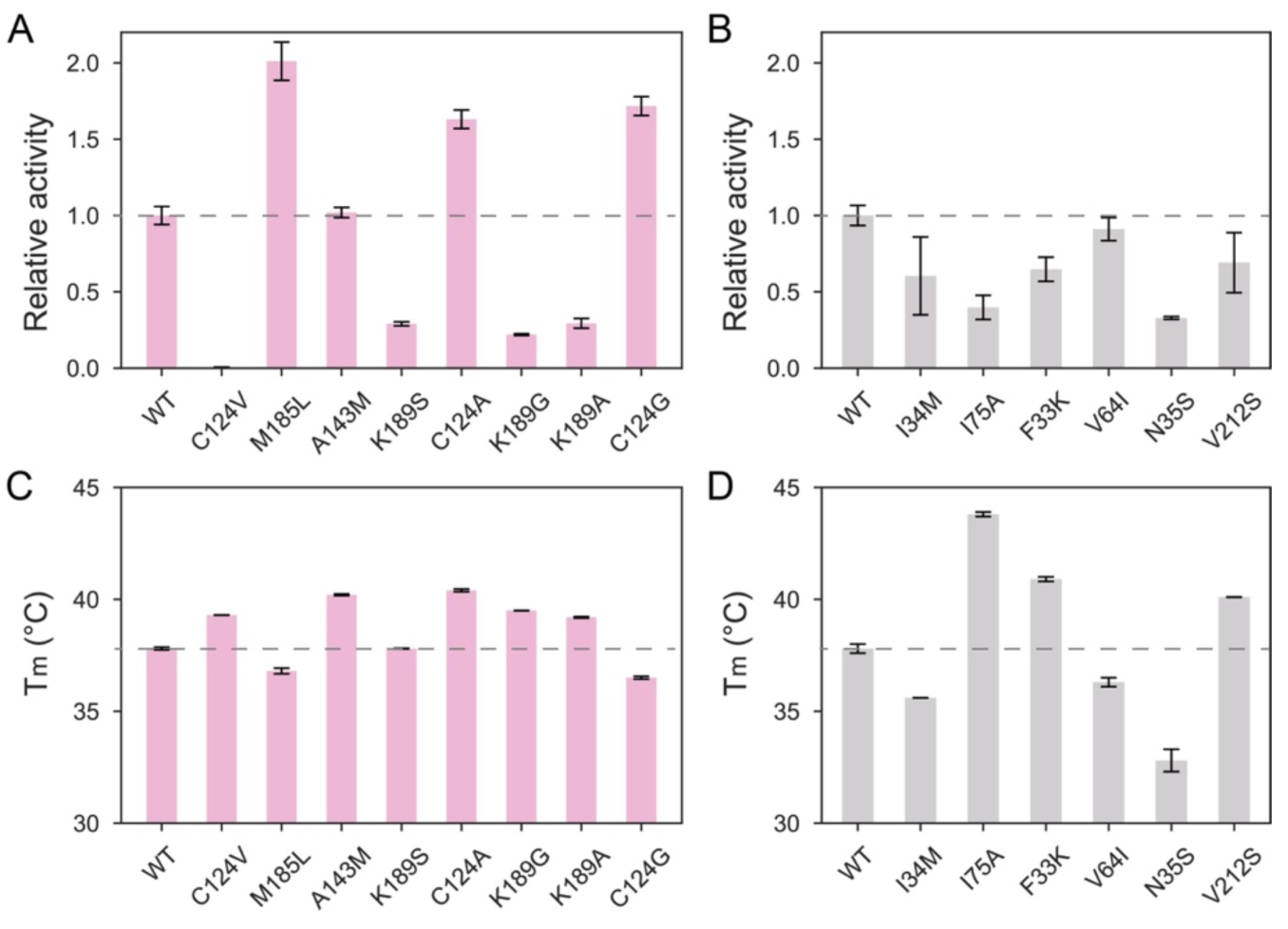
Activity and stability of designed variants in the experiment. (A, B) Relative activity for variants mutated in the active center (A) and on the enzyme scaffold (B). (C, D) Melting temperature for variants mutated in the active center (C) and on the enzyme scaffold (D). Values of relative activity and melting temperature used in the plot were given in Table S6 and S7, respectively.

It is noteworthy that the C124A mutation was previously identified as a potential target for mutation due to concerns that the cysteine residue may be prone to oxidation and impair enzyme activity.^39^ This mutation was subsequently used in different luciferase engineering projects.^32 33^ Our approach also identifies C124G as a beneficial mutation. However, it would be valuable to further investigate the chemical mechanisms underlying the observed effects, as C124V, another mutation on the same residue results in an inactive luciferase. Our method also suggested that the M185L mutation could improve catalysis. The methionine residue at residue 185, located close to the substrate, may be able to adjust its orientation to enhance activity. This is in line with previous findings, which identified a beneficial valine mutation in this location.^32^ However, we found that three proposed K189 mutations were not effective in practice. Enzyme design is indeed a challenging field, and it is extremely hard to understand why certain designs fail.^15 16 17 18 19^ Additional investigations are necessary to comprehend the molecular intricacies underlying results present here, which extends beyond the scope of the generative AI studies conducted in this work.

For mutations on the enzyme scaffold, all six expressed variants show slightly decreased activity (Fig 4B). The discovery is fascinating and aligns with our earlier conclusion that the primary evolutionary driving force for the enzyme scaffold is not centered on improving enzyme efficiency. ^22^ Changes to the enzyme scaffold have a less pronounced effect on activity than changes in the active center, further supporting the idea that enzyme activity drives the evolution of residues near the substrate. We also analyzed the emission spectra for each design (Fig S8). The shift in the spectrum caused by mutations to the active center is more evident compared to those on the enzyme scaffold.

Together, our computational and experimental results provide strong support for using natural evolutionary information to improve activity by focusing on the active center. However, this does not rule out the potential for discovering advantageous mutations in other regions using evolutionary data.^27^

### Enhancing Luciferase Stability on the Enzyme Scaffold Validated in Experiments

The stability of an enzyme is another crucial property. We have previously observed that the enzyme scaffold is primarily shaped by evolutionary pressure to maintain stability for some enzymes.^22^ It will be a crucial test of the evolution-stability correlation by measuring the melting temperature of variants designed here.

Our method primarily recommends mutations in the active center to enhance biological activity, not stability.^22^ Therefore, it is not expected that these mutations will systematically improve enzyme stability. We measured the stability of eight variants in the active center using differential scanning fluorimetry assay (Fig S9). In line with our expectations, compared to the wild-type enzyme, they all have similar melting temperatures (*T*_m_) (Fig 4C) and the change in *T*_m_ ranges from -1.3 °C to 2.4 °C.

Conversely, mutations on the scaffold of the enzyme can significantly affect the stability of RLuc, causing *T*_m_ to vary between -5.0 °C and 6.0 °C (Fig 4D). Our method successfully identified three out of six mutations that led to an increase in stability, resulting in a 50% success rate. The I75A mutant alone can increase the melting temperature by 6.0 °C. These stability results suggest that natural evolutionary information can be utilized to design stabilizing mutations away from the active regions, again aligning with the conclusions presented in Ref ^22^.

### Comparison of Enzyme Engineering Efforts Utilizing Generative Models

Generative models, trained on natural homologous sequences, have shown potential in generating functional sequences akin to those found in nature.^15 16 17 18^ As outlined in Table 1, recent efforts have employed various generative models capable of producing functional enzyme sequences. Despite these advancements, engineering enzymes to exceed the performance of their wild-type counterparts remains a considerable challenge.

**Table 1.** Activity of designed variants using generative AI.

| Enzyme | Generative Model | Mutant Type | Functional Assay Comment | Activity Assay Comment |
| --- | --- | --- | --- | --- |
| Luciferase RLuc | MaxEnt (this study) | Single |  | 4 out of 8 designs show enhanced activity compared to wild-type; ***maximum fold increase: 2.0*** |
| Chorismate Mutase | MaxEnt <sup>15</sup> | Multiple | 481 out of 1618 are functional | 5 tested designs exhibit comparable activity to natural enzymes |
| Malate Dehydrogenase | GAN <sup>16</sup> | Multiple | 13 out of 55 are functional |  |
| Luciferase LuxA | VAE <sup>17</sup> | Multiple | 9 out of 11 are functional |  |
| Lysozyme | Language <sup>18</sup> | Multiple |  | 6 tested designs exhibit comparable activity compared to wild-type |
| Ornithine Transcarbamylase | VAE <sup>19</sup> | Multiple |  | 58 out of 87 designs show enhanced activity compared to wild-type; ***maximum fold increase: 2.5*** |

\*GAN: Generative Adversarial Network; VAE: Variational Autoencoder; Language: Language Model

On a different note, in Ref ^19^, VAE was employed to facilitate the design of variants with enhanced activity through the incorporation of multiple mutations, realizing a maximum fold increase of 2.5. Notably, our approach enabled us to introduce a single mutation, leading to a fold increase of 2.0, a result that is on par with those achieved by the introduction of multiple mutations. In practical scenarios, fewer mutations are favorable as they minimize experimental effort and potential ethical issues.^20^ Another distinction is that we can enhance the activity by mutating residues near the substrate, which is not the case in Ref ^19^. It can be attributed to our establishment of the evolution-catalysis relationship using generative models, which provides clues to effectively harness evolutionary information for rational enzyme engineering.

## Discussions

This study reinforces our previously proposed connection between evolution and catalysis. Our suggested mutations, derived from sequences evolved in nature, successfully enhance enzyme activity in the active center and bolster stability on the protein scaffold. The engineering success underscores the potential of generative AI to design enzymes. Nonetheless, we did not achieve a flawless success rate in designing beneficial single variants. This could be due to limitations in classifying residues solely based on their distance to the substrate, an incomplete categorization of the enzyme architecture, a current lack of understanding of the complexities of luciferase evolution, etc. Beyond single variants, it is essential to examine the maximal fold increase by assessing higher-order variants experimentally. Combining advantageous single mutations with other neutral ones could potentially produce many improved diverse variants, potentially adding complexity to pinpointing the optimal higher-order ones. It remains to be seen whether this applies to Ref ^19^, especially since its fold increase mirrors that of a single mutant.

The MaxEnt model is based on the construction of protein MSA. Meanwhile, the protein language model, trained on millions of natural sequences, might discern common protein patterns. Like the MaxEnt model, this language model is generative. A question then arises: how well would a language model elucidate the evolution-catalysis relationship for luciferase? To this end, we utilized ESM-1v,^40^ trained on 98 million natural sequences, to rank luciferase variants with mutations proximate to the substrate. The correlation derived was somewhat inferior to the MaxEnt model that leverages an MSA (Fig S10). Several explanations are plausible. Firstly, the variants are predominantly double mutants; the scoring function in ESM-1v treats mutated positions in isolation, overlooking epistasis. Secondly, the myriad of enzyme-catalyzed reactions suggests diverse mechanisms; extracting universally applicable rules might be challenging. Hence, interspersing information across different protein classes might not bolster the prediction of enzyme activity. However, to definitively ascertain whether a protein language model surpasses the MSA-based generative model’s efficacy, comprehensive evaluations across larger datasets are imperative.

We now turn our attention to the connection between designs derived from generative models and consensus design. In consensus design, the most frequent residue type within a protein family is selected.^41,42^ We showcased the sequence logo for the mutated positions in our experiment in Fig S11, emphasizing the conservation of sequences at these positions. Some single variants proposed by the MaxEnt model align with the consensus residue at specific positions. For instance, for mutations within the active center, A143M selects the consensus methionine. Yet, the consensus residues at positions 124, 185, and 189 are not the best choices as per the MaxEnt model. Concerning mutations on the scaffold, five designs, namely I34M, I75A, V64I, Y298A, and V212S, align with consensus design. This consistency is anticipated since a MaxEnt model, when not accounting for epistasis, independently evaluates each position using the statistical energy 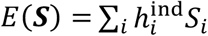 and tends to favor the consensus residue. Yet, the evolution-catalysis relationship provides rationale for why consensus design might be effective. Furthermore, the MaxEnt model is anticipated to surpass the consensus design when considering higher-order mutants.

The brighter luciferase could have applications in single-cell imaging, and evolutionary constraints have driven the evolution of blue light in RLuc, whereas red-shifted light is optimal for deep tissue penetration. It is notable that modifications to the substrate, rather than enzyme engineering, have largely been adopted to tune luciferase spectrum. Additionally, luciferase has evolved independently multiple times on Earth, such as in fireflies that could produce fluorescence with red light which is different from RLuc. The specific reasons for these different evolutionary trends remains a mystery, and further effort is needed to better understand the factors that drive the evolution of bioluminescence and luciferase activity in different organisms.

The remarkable catalytic properties of enzymes evolved by nature serve as a valuable source of inspiration for protein engineering. However, there is still much to be learned about the underlying principles of natural evolution that can inform rational enzyme engineering. Our study contributes to the advancement of this understanding and provides new insights into rational enzyme design.

## Supporting information

RLuc MSA

## Acknowledgment

A.W. is supported by the National Institutes of Health R35 GM122472 and the National Science Foundation Grant MCB 1707167. S.D. is supported by the National Natural Science Foundation of China (Nos. 22177004 and 92153301). W.J.X. is supported by the startup funding from University of Florida. We thank the High Performance Computing and Communication Center at the University of Southern California and the Hypergator at the University of Florida for providing computational resources. We thank Tianmin Fu, Zhenghan Liao, and Xiaoyu Chen for their insightful discussions.

## Additional information

Supplementary information and dataset are available for this paper.

## Data Availability

The RLuc MSA are uploaded as supporting files. The calculated statistical energies for mutants are reported in the supplementary information.

