## supplemental information for "Enhancing Luciferase Activity and Stability through Generative Modeling of Natural Enzyme Sequences"

### Supporting Information

#### Embedding protein sequences in low-dimensional latent space using VAE

VAE was adopted to perform the low-dimensional embedding. The one-hot representation of the sequence data $\boldsymbol{S}$ was projected to the latent space $\boldsymbol{z}$. $q(\boldsymbol{z}|\boldsymbol{S})$ and $p(\boldsymbol{S}|\boldsymbol{z})$ are the neural networks that encode the sequence data to latent variables and decode from latent space to reconstruct the original sequence data, respectively. To train the model, we optimized the evidence lower bound $E_q[\log p(\boldsymbol{S}|\boldsymbol{z})] - D_{KL}[q(\boldsymbol{z}|\boldsymbol{S})||p(\boldsymbol{z})]$, where $p(\boldsymbol{z})$ is the Gaussian prior of the latent variables, KL indicates Kullback-Leibler divergence.

The VAE model was implemented in PyTorch and the Adam optimizer was selected to optimize the model. One hidden layer with 100 nodes was used for the encoding and decoding neural network. For visualization, a two-dimensional latent space was chosen.

#### Construction of RLuc variants and protein production

To construct a bacterial expression plasmid, the wild-type RLuc luciferase and its variants were cloned into pET21b vector with optimized sequence for E. coli and 6xHis tag for protein purification. As for the preparation of plasmids for different variants, QuickMutation™Plus Site-Directed Mutagenesis Kit (Beyotime, D0208S) was utilized to introduce site-specific mutations following manufacture instructions. The luciferase expression plasmids were transformed into DH5α competent cells (Solarbio, C1100) for amplification, followed by verifying the sequence with Sanger sequencing. As for the expression of wild-type luciferase and its site-specific variants, the plasmids were transformed into BL21-competent cells (Solarbio, C1410). Expression of the corresponding genes was conducted in 100 ml LB medium (10 ug/ml ampicillin) at 220 rpm and 37 °C, followed by adding 0.5 mM isopropyl β-D-1-thiogalactopyranoside (IPTG) when the $OD_{600}$ value reached 0.6-0.8. Then, the cells were cultured for 16-20 h at 220 rpm and 16 °C for protein expression.

The cells were centrifuged at 4000 rpm for 40 min at 4 °C, and the supernatant was discarded. Then, the cell pellet was resuspended with 20 ml of lysis buffer (0.1 M Tris, 0.1% TritonX-100, pH = 8) and treated with sonication for 30 min at 4 °C to enable total release of the protein from the cells. The mixture was centrifuged at 12000 rpm for 40 min and 4 °C, then filtered the supernatant with a 0.2 um filter. The crude was purified with a nickel affinity column (HisTrap HP, 5 ml). Firstly, the column was balanced with binding buffer (50 mM Tris, 5 mM imidazole, and 500 mM NaCl) before the crude was loaded on the column. 20% of elution buffer (50 mM Tris, 200 mM imidazole, and 500 mM NaCl) was utilized to wash the column before eluting with 100% of elution buffer to provide target proteins. The collected wild-type luciferase and variants were quantified with a BCA protein quantification kit (Bioss, C05-02001), characterized with SDS-PAGE, and stored at 4 °C before use.

**Measurement of spectrum**

The spectrum of wild-type luciferase and its different variants was measured with FlexStation 3 Multi-Mode Microplate Reader. Specifically, 10 µl of coelenterazine working solution was added to 100 µl of luciferase variants (50 ng/ml in 0.1 M sodium phosphate buffer pH 7.0), followed by reading the luminescence intensity at the range of wavelength from 300 nm to 600 nm (step length is 1 nm and integration time is 10 ms).

**Measurement of activity**

Coelenterazine (from TargetMol) was dissolved with propanediol at a concentration of 2 mg/ml as a stock solution. The luciferase and its variants were diluted with sodium phosphate buffer (0.1 M, pH 7.0) at the final concentration of 0.5 ng/ml, and 100 µl of each mutant was placed into the 96-well plate with five replicates. Coelenterazine was diluted to 0.2 ug/ml with sodium phosphate buffer as a working solution. Luciferase activity was measured by adding 10 µl of coelenterazine working solution to each well at room temperature, shaking for 3 s, delaying for 3 s, and reading for 10 s with Centro LB 960 microplate luminometer sequentially. Despite exhibiting discernible luminescence, C124V has very low activity. The emission spectrum remains largely unaffected by active variants, thus any variance in luminescence intensity can be primarily attributed to differences in enzyme activity. Therefore, the relative activity of different variants was calculated by normalizing the luminescence intensity of wild-type luciferase.

**Measurement of stability**

As for the stability evaluation, wild-type luciferase and its variants (0.5 ng/ml in sodium phosphate buffer) were incubated with different times at 30 °C respectively, transferred to 4 °C for more than 5 min and measured for the activity following the procedure described above. The melting temperature of each mutant was determined by differential scanning fluorimetry assay using Prometheus NT.48., from 20 °C to 90 °C.

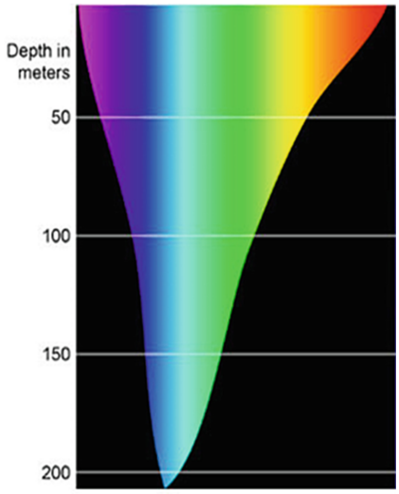


**Figure S1**. Spectral attenuation of visible light wavelengths (400 nm - 700 nm) in open water ( https://commons.wikimedia.org/wiki/File:NOAA_Deep_Light_diagram3.jpg ).

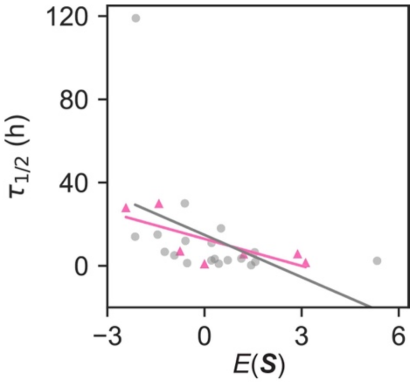


**Figure S2**. Correlations between the statistical energy $E(\boldsymbol{S})$ and inactivation time measured in the mouse. The correlation values for the active center and enzyme scaffold are -0.74 ($p$-value = 0.054) and -0.43 ($p$-value = 0.066), respectively. The correlation values are similar to those measured in the rat. Correlations were calculated using data in Table S1.

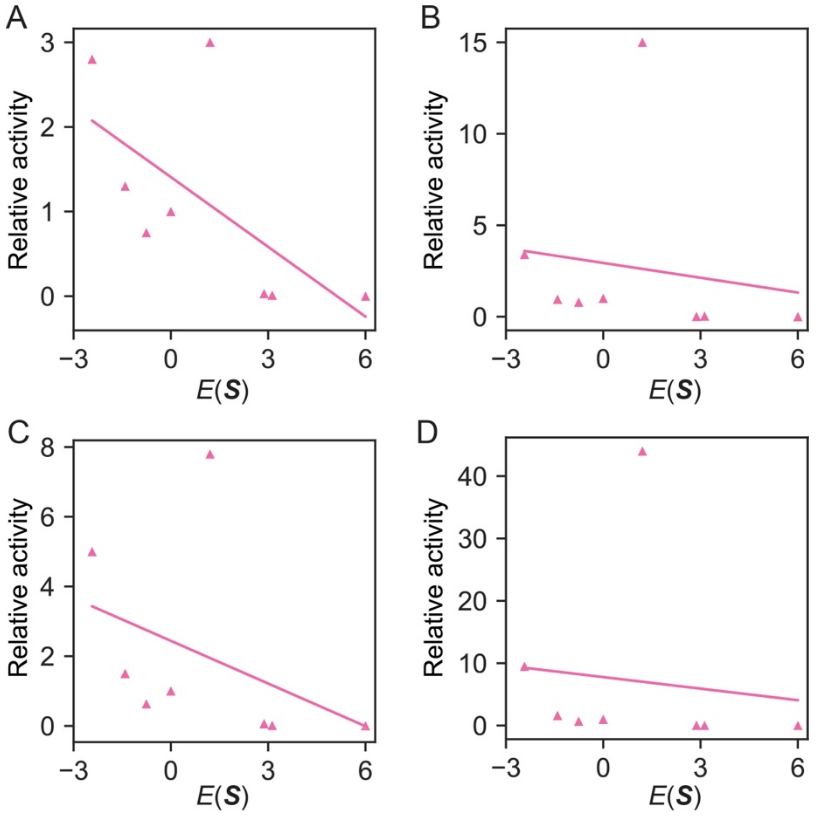


**Figure S3**. Correlations between the statistical energy $E(\boldsymbol{S})$ and luciferase activity using different substrates. (A) Benzy-coelenterazine (bc): correlation value of -0.64 ($p$-value = 0.090). (B) Coelenterazine-cp (cp): correlation value of -0.15 ($p$-value = 0.73). (C) Coelenterazine-n (n): correlation value of -0.40 ($p$-value = 0.33). (D) Bisdeoxycoelenterazine (bdc): correlation value of -0.11 ($p$-value = 0.79). Correlations were calculated using data in Table S1.

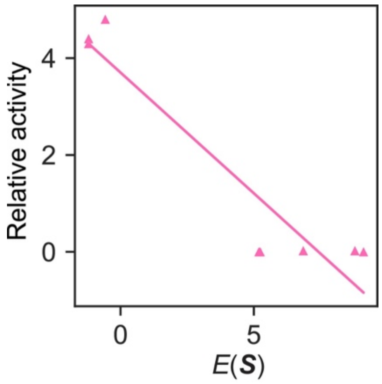


**Figure S4**. Correlation between the statistical energy $E(\boldsymbol{S})$ and luciferase catalysis for RLuc8 variants: correlation value of -0.94 ($p$-value < 0.001). The correlation was calculated using data in Table S2.

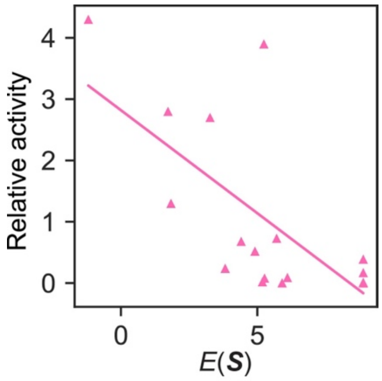


**Figure S5**. Correlation between the statistical energy $E(\boldsymbol{S})$ and luciferase catalysis for RLuc8 variants with mutations on residues I233: correlation value of -0.71 ($p$-value < 0.001). The correlation was calculated using data in Table S3.

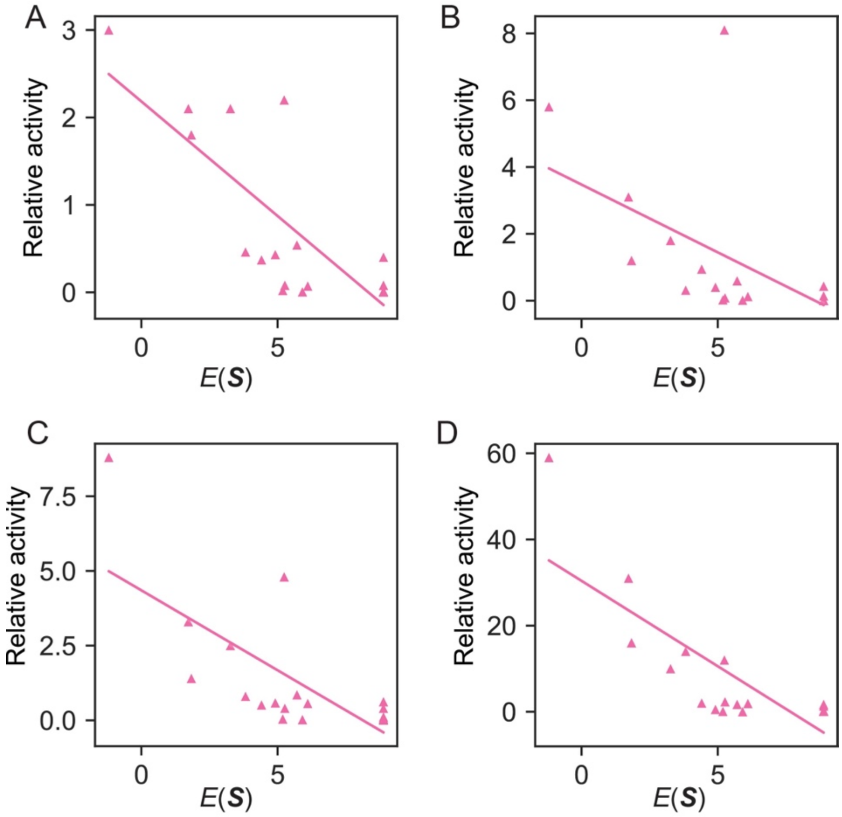


**Figure S6**. Correlations between the statistical energy $E(\boldsymbol{S})$ and luciferase catalysis for RLuc8 variants with mutations on residues I233 using different substrates. (A) Benzy-coelenterazine (bc): correlation value of -0.79 ($p$-value < 0.001). (B) Coelenterazine-cp (cp): correlation value of -0.55 ($p$-value = 0.012). (C) Coelenterazine-n (n): correlation value of -0.72 ($p$-value < 0.001). (D) Bisdeoxycoelenterazine (bdc): correlation value of -0.80 ($p$-value < 0.001). Correlations were calculated using data in Table S3.

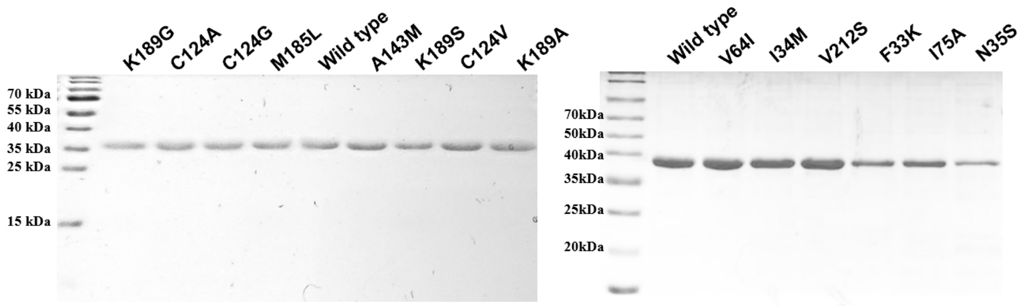


**Figure S7**. Characterization of RLuc and its variants with SDS-PAGE

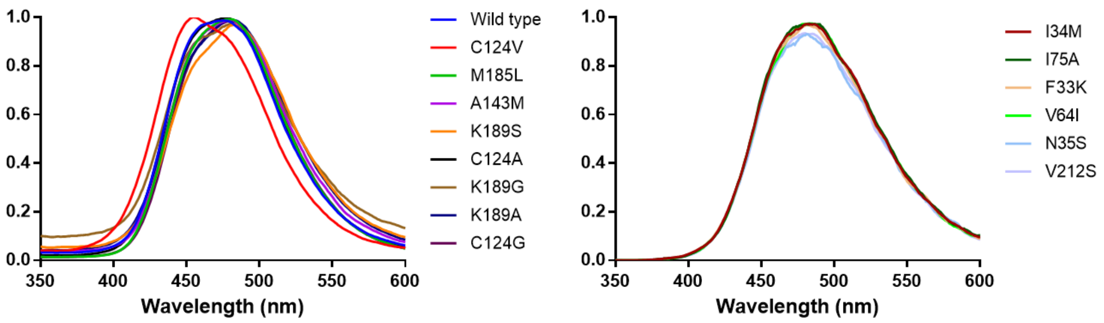


**Figure S8**. Normalized bioluminescence emission spectra of RLuc and its variants

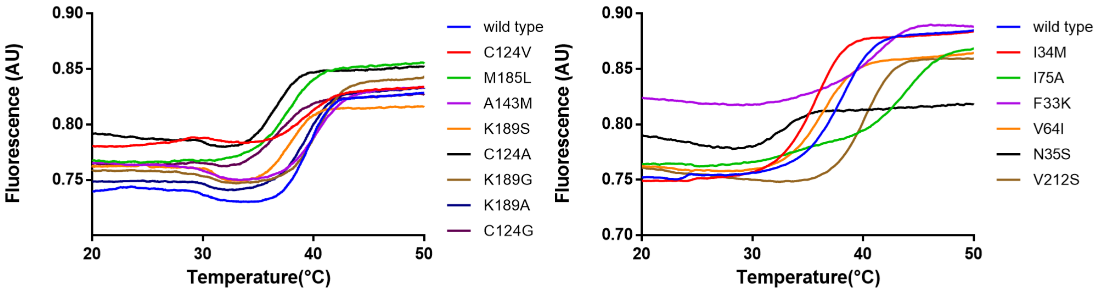


**Figure S9**. DSF melting curves of RLuc and its variants.

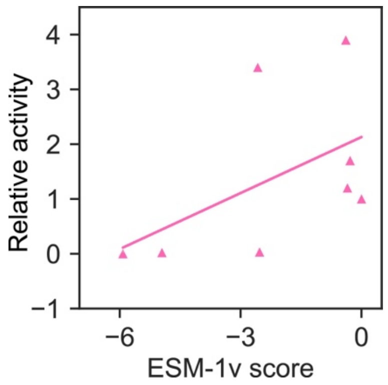


**Figure S10**. Relationship between ESM-1v scores and luciferase activity within the active center. The correlation coefficient stands at 0.51, with a p-value of 0.15. The ESM-1v score is formulated as the logarithmic odds ratio comparing a mutated variant to the wild-type protein, as determined by a pretrained protein language model. This analysis was conducted using the ESM model available at https://github.com/facebookresearch/esm , with results averaged across five pretrained models.

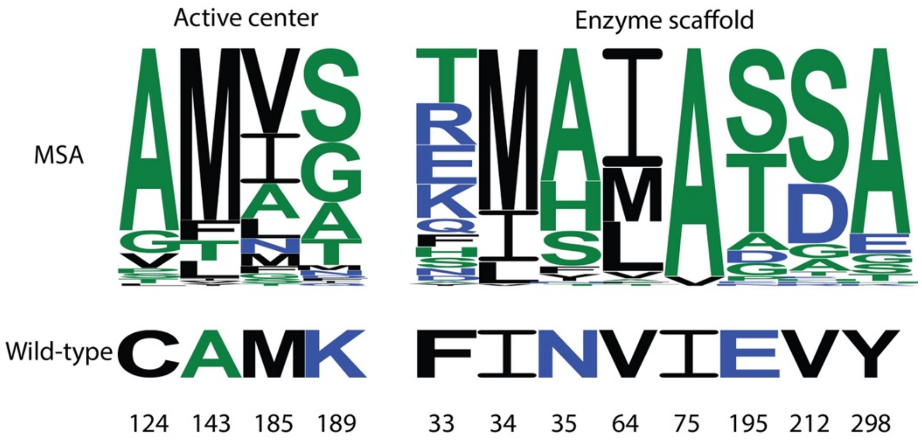


**Figure S11**. Sequence logo representing the positions mutated during the experiment. Within the active center, mutations occurred at four positions, while the enzyme scaffold had mutations at eight distinct positions.

**Table S1**. Mutations of RLuc in previous luciferase engineering experiments

| | Relative activity | | | | | Inactivation $\tau_{1/2}$ (h) | | Wavelength (nm) | | | |
|---|---|---|---|---|---|---|---|---|---|---|---|
| mutant | Native | bc | cp | n | bdc | Mouse | Rat | Peak | Mean | d (Å) | $E(\boldsymbol{S})$ |
| Wild-type | 1 | 1 | 1 | 1 | 1 | 1 | 1 | 482 | 497 | 0 | 0 |
| C124A | 1.2 | 0.75 | 0.79 | 0.63 | 0.68 | 7.1 | 6.6 | 482 | 498 | 7.91 | -0.76 |
| F33R/I34M/C124A | 0.15 | 0.15 | 0.16 | 0.12 | 0.2 | 0.3 | 0.3 | 481 | 497 | 17.19 | 1.44 |
| E44G/C124A | 0.94 | 0.78 | 0.74 | 0.66 | 0.98 | 2.6 | 3.3 | 486 | 502 | 18.60 | 0.21 |
| A54G/A55G/C124A | 0.12 | 0.1 | 0.06 | 0.15 | 0.19 | 2.4 | 3 | 476 | 492 | 9.33 | 5.33 |
| A54P/A55T/C124A | 0.21 | 0.15 | 0.11 | 0.38 | 0.22 | 119 | 129 | 470 | 483 | 9.33 | -2.12 |
| A54P/C124A | 0.05 | 0.04 | 0.05 | 0.08 | 0.06 | 14 | 13 | 468 | 482 | 8.76 | -2.14 |
| A55T/C124A | 1.7 | 1.2 | 0.58 | 1.4 | 2.4 | 30 | 29 | 486 | 504 | 9.20 | -0.61 |
| F116L/C124A | 1.3 | 1 | 1.3 | 0.88 | 1.8 | 11 | 9.4 | 486 | 502 | 11.69 | 0.22 |
| C124A/S130A | 1.7 | 1.4 | 1.7 | 1.4 | 2.6 | 18 | 14 | 482 | 498 | 12.05 | 0.51 |
| C124A/K136R | 2.5 | 2.1 | 1.9 | 1.9 | 2.6 | 12 | 11 | 482 | 498 | 15.38 | -0.59 |
| C124A/A143M | 1.7 | 1.3 | 0.95 | 1.5 | 1.6 | 30 | 29 | 480 | 497 | 7.10 | -1.41 |
| C124A/F180A | 0.02 | 0.01 | 0.03 | 0.01 | 0.01 | 1.6 | 1.6 | 488 | 504 | 6.37 | 3.12 |
| C124A/M185V | 3.4 | 3 | 15 | 7.8 | 44 | 5.7 | 3.7 | 485 | 500 | 6.24 | 1.21 |
| C124A/M191L | 1.1 | 0.99 | 0.97 | 1 | 1.2 | 6.5 | 5.1 | 480 | 496 | 11.37 | 1.55 |
| C124A/E195S/P196D | 0.12 | 0.1 | 0.12 | 0.1 | 0.15 | 1 | 0.7 | 482 | 498 | 16.96 | 0.44 |
| C124A/F199M | 0.58 | 0.44 | 0.53 | 0.49 | 0.46 | 6.7 | 6 | 480 | 495 | 11.40 | -1.23 |
| C124A/L203R | 0.55 | 0.55 | 0.52 | 0.41 | 0.43 | 2.7 | 2.2 | 484 | 501 | 12.65 | 0.72 |
| C124A/G229E | 0.02 | 0.01 | 0.03 | 0.03 | 0.01 | 1.9 | 1.8 | 473 | 490 | 8.99 | 1.57 |
| C124A/Q235A | 1.2 | 1.1 | 1.1 | 1 | 1.2 | 3.3 | 3.6 | 473 | 489 | 10.12 | 0.31 |
| C124A/M253L | 1.9 | 1.4 | 1.6 | 1.6 | 1.7 | 15 | 10 | 471 | 488 | 10.84 | -1.45 |
| C124A/S257G | 1.1 | 0.95 | 1.3 | 1.1 | 3 | 1.3 | 1.4 | 477 | 493 | 8.77 | -0.53 |
| C124A/F261L/F262L | 0 | 0 | 0 | 0 | 0 | | | | | 5.06 | 6.00 |
| C124A/F262L | 0.03 | 0.03 | 0.01 | 0.06 | 0.03 | 5.8 | 6.4 | 478 | 495 | 5.74 | 2.88 |
| C124A/S287L | 3.9 | 2.8 | 3.4 | 5 | 9.5 | 28 | 20 | 478 | 496 | 8.48 | -2.43 |
| C124A/M295I | 1 | 0.83 | 0.57 | 0.72 | 0.86 | 5 | 4.9 | 480 | 497 | 10.62 | -0.93 |
| C124A/K300A | 1.1 | 1 | 1.1 | 1 | 1.3 | 3.5 | 3.9 | 481 | 497 | 15.45 | 1.14 |

*The experimental values were obtained from *Protein Engineering, Design and Selection* **19**, 391–400 (2006), including relative activity, inactivation time, and wavelength. The average distance (d) between the substrate and mutated residues was calculated using the crystal structure with PDB ID: 6YN2.

**Table S2**. Mutations of RLuc8 in previous luciferase engineering experiments

| | Relative activity | | | | | Inactivation $\tau_{1/2}$ (h) | | Wavelength (nm) | | | |
|---|---|---|---|---|---|---|---|---|---|---|---|
| mutant | Native | bc | cp | n | bdc | Mouse | Rat | Peak | Mean | d (Å) | $E(\boldsymbol{S})$ |
| RLuc8 | 4.3 | 3 | 5.8 | 8.8 | 59 | 281 | 86 | 487 | 503 | 11.39 | -1.20 |
| D120A | 0 | 0 | 0.001 | 0 | 0.21 | | | | | 10.51 | 5.24 |
| D120N | 0.02 | 0.02 | 0.05 | 0.34 | 5.1 | | | | | 10.51 | 6.86 |
| E144A | 0 | 0 | 0 | 0 | 0 | 57 | 13 | | | 10.49 | 9.12 |
| E144Q | 0 | 0 | 0 | 0 | 0.002 | | | | | 10.49 | 5.19 |
| H285A | 0.02 | 0.02 | 0.046 | 0.03 | 0.2 | | 21 | | | 10.51 | 8.79 |
| M185V | 4.4 | 2.6 | 12 | 4.1 | 20 | 0.8 | 0.3 | | | 10.63 | -1.20 |
| M185V/Q235A | 4.8 | 2.7 | 14 | 7.1 | 20 | 0.5 | 0.2 | | | 10.80 | -0.57 |

*The experimental values were obtained from *Protein Engineering, Design and Selection* **19**, 391–400 (2006), including relative activity, inactivation time, and wavelength. The average distance (d) between the substrate and mutated residues was calculated using the crystal structure with PDB ID: 6YN2.

**Table S3**. Mutations of RLuc8 on the I233 residue in previous luciferase engineering experiments

| | Relative activity | | | | | |
|---|---|---|---|---|---|---|
| mutant | Native | bc | cp | n | bdc | $E(\boldsymbol{S})$ |
| RLuc8 | 4.3 | 3 | 5.8 | 8.8 | 59 | -1.20 |
| I223A | 0.68 | 0.37 | 0.94 | 0.51 | 2 | 4.41 |
| I223C | 3.9 | 2.2 | 8.1 | 4.8 | 12 | 5.24 |
| I223D | 0.01 | 0.01 | 0.01 | 0.06 | 0.1 | 8.88 |
| I223E | 0.01 | 0.01 | 0.01 | 0.11 | 0.21 | 8.88 |
| I223F | 2.7 | 2.1 | 1.8 | 2.5 | 10 | 3.27 |
| I223G | 0.17 | 0.08 | 0.14 | 0.4 | 1.3 | 8.88 |
| I223H | 0.09 | 0.07 | 0.12 | 0.56 | 1.9 | 6.10 |
| I223K | 0.002 | 0.002 | 0.001 | 0.003 | 0.26 | 8.88 |
| I223L | 1.3 | 1.8 | 1.2 | 1.4 | 16 | 1.83 |
| I223M | 0.24 | 0.46 | 0.31 | 0.8 | 14 | 3.82 |
| I223N | 0.39 | 0.4 | 0.43 | 0.62 | 1.7 | 8.88 |
| I223P | 0.01 | 0.01 | 0.01 | 0.03 | 0.13 | 8.88 |
| I223Q | 0.08 | 0.08 | 0.08 | 0.4 | 2.3 | 5.26 |
| I223R | 0.004 | 0.003 | 0.003 | 0.01 | 0.24 | 8.88 |
| I223S | 0.73 | 0.54 | 0.59 | 0.85 | 1.7 | 5.71 |
| I223T | 0.52 | 0.43 | 0.4 | 0.58 | 0.54 | 4.91 |
| I223V | 2.8 | 2.1 | 3.1 | 3.3 | 31 | 1.72 |
| I223W | 0.003 | 0.004 | 0.01 | 0.02 | 0.01 | 5.91 |
| I223Y | 0.02 | 0.02 | 0.02 | 0.04 | 0.07 | 5.19 |

*The experimental values were obtained from *Nature Methods* **4**, 641-643 (2007).

**Table S4**. Variants with redesigned active center from the MaxEnt model

| mutant | $E(\boldsymbol{S})$ | mutant | $E(\boldsymbol{S})$ | mutant | $E(\boldsymbol{S})$ |
|---|---|---|---|---|---|
| Wild-type | 0.00 | K189A/N264E | -0.31 | A123S/K189S | -0.67 |
| S188N/K189A | 0.00 | C124G/N264D | -0.31 | C124G/N241S | -0.68 |
| A143M/G269W | 0.00 | K189A/N241S | -0.32 | K189A/G269I | -0.68 |
| V149P/K189A | -0.02 | A123S/M185L | -0.32 | D148K/K189G | -0.69 |
| C124G/E268D | -0.02 | L165A/K189A | -0.32 | C124V/A143M | -0.70 |
| C124S/K189G | -0.02 | C124G/N264A | -0.32 | D158E/K189G | -0.71 |
| C124G/G269R | -0.03 | C124A/K189D | -0.32 | D148S/K189A | -0.72 |
| K189G/N264E | -0.03 | C124A/D148A | -0.33 | C124G/G269T | -0.73 |
| K189A/G269Y | -0.04 | T184R/K189S | -0.33 | K189G/F273W | -0.73 |
| K189A/G269A | -0.04 | C124G/V237I | -0.33 | K189S | -0.73 |
| C124G/W156L | -0.05 | C124P/K189G | -0.33 | K189G/G269V | -0.73 |
| I150L/K189G | -0.05 | A143F/K189G | -0.34 | A123S/K189G | -0.75 |
| D148A/M185L | -0.05 | C124G/K189I | -0.34 | C124A | -0.76 |
| C124G/I166M | -0.05 | C124T/M185L | -0.34 | D148A/K189A | -0.78 |
| C124A/I166L | -0.05 | K189A/L225F | -0.34 | C124G/N264E | -0.80 |
| A143M/L165A | -0.06 | C124G/N264G | -0.35 | C124V/K189S | -0.80 |
| A143M/I166L | -0.07 | D148S/K189G | -0.35 | C124G/N264P | -0.82 |
| C124G/K189H | -0.07 | C124G/G269L | -0.36 | C124G/I150L | -0.83 |
| C124G/E268K | -0.07 | A143M/K189N | -0.36 | C124G/K189Q | -0.85 |
| A143F/K189S | -0.07 | K189G/G269I | -0.36 | C124G/L165A | -0.85 |
| I166L/K189S | -0.08 | A143M/D148R | -0.37 | C124G/W156F | -0.86 |
| I150L/K189S | -0.08 | K189A/N264P | -0.37 | C124T/K189G | -0.86 |
| K189S/N264K | -0.08 | C124G/S188N | -0.37 | C124G/K189F | -0.87 |
| C124P/A143M | -0.08 | C124G/I150M | -0.38 | K189G | -0.87 |
| A143M/G269F | -0.09 | C124G/A143T | -0.38 | C124G/L225F | -0.96 |
| T184R/K189G | -0.09 | C124G/V149P | -0.38 | D158E/K189A | -0.96 |
| C124I/K189G | -0.10 | C124S/K189A | -0.39 | D148R/K189A | -0.98 |
| C124V | -0.10 | M185L | -0.40 | A143M/M185L | -1.00 |
| D148G/K189A | -0.11 | T184R/K189A | -0.40 | C124G/T184R | -1.02 |
| C124A/A143F | -0.11 | C124G/G269E | -0.41 | K189A/G269V | -1.04 |
| A143M/D148S | -0.11 | C124I/K189A | -0.41 | C124G/I166L | -1.05 |
| C124A/I150L | -0.11 | A143M/D148K | -0.41 | C124G/G269F | -1.05 |
| A143M/G269I | -0.12 | A143M/K189M | -0.43 | C124G/N264K | -1.06 |
| C124A/N264K | -0.14 | I150L/K189A | -0.44 | C124G/A143F | -1.07 |

| | | | | | |
|---|---|---|---|---|---|
| D148K/M185L | -0.15 | C124G/K189V | -0.45 | A123S/K189A | -1.07 |
| K189G/L225F | -0.15 | C124V/M185L | -0.48 | D148K/K189A | -1.08 |
| A143M/K189- | -0.16 | D148R/K189S | -0.49 | C124G/G269W | -1.10 |
| K189S/G269W | -0.16 | C124G/K189R | -0.50 | C124V/K189G | -1.11 |
| A143M/K189D | -0.16 | A143M/G269V | -0.51 | K189A/F273W | -1.12 |
| D148S/K189S | -0.17 | C124G/G269A | -0.52 | M185L/K189S | -1.13 |
| D148Q/K189A | -0.17 | C124A/K189N | -0.52 | C124G/D148S | -1.15 |
| D158E/M185L | -0.17 | I166L/K189A | -0.53 | C124G/G269I | -1.16 |
| A143M/D148A | -0.17 | D148A/K189G | -0.54 | C124T/K189A | -1.18 |
| C124P/K189S | -0.18 | C124G/G269Y | -0.54 | C124G/K189D | -1.20 |
| M185L/K189N | -0.18 | A143M/D158E | -0.54 | K189A | -1.21 |
| M185L/K189M | -0.18 | C124A/K189M | -0.55 | C124G/K189- | -1.22 |
| C124G/I159S | -0.18 | K189A/N264K | -0.55 | C124A/M185L | -1.25 |
| I166L/K189G | -0.18 | D148K/K189S | -0.55 | M185L/K189G | -1.28 |
| C124A/T184R | -0.20 | A123S/A143M | -0.55 | C124G/D148A | -1.29 |
| C124A/D148S | -0.21 | D158E/K189S | -0.56 | C124V/K189A | -1.33 |
| K189S/G269I | -0.21 | C124G/K189W | -0.57 | C124A/A143M | -1.41 |
| K189A/L225I | -0.22 | A143M/K189T | -0.57 | C124G/D148R | -1.42 |
| C124A/G269F | -0.22 | A143M/F273W | -0.58 | C124G/K189N | -1.44 |
| D148R/M185L | -0.23 | A143F/K189A | -0.58 | A143M/K189S | -1.45 |
| C124G/N241A | -0.23 | C124T/A143M | -0.58 | C124G/D148K | -1.49 |
| C124G/D148M | -0.24 | C124G/D148G | -0.58 | A143M/K189G | -1.52 |
| K189S/G269F | -0.24 | C124A/D158E | -0.59 | C124G/K189M | -1.53 |
| K189G/G269F | -0.24 | C124G/D148Q | -0.59 | C124G/G269V | -1.58 |
| K189A/G269T | -0.24 | C124A/G269V | -0.59 | C124G/F273W | -1.59 |
| C124G/L225V | -0.25 | C124A/F273W | -0.60 | C124G/D158E | -1.59 |
| D148A/K189S | -0.25 | K189S/G269V | -0.60 | C124A/K189G | -1.61 |
| C124A/G269I | -0.26 | K189A/G269F | -0.60 | C124G/K189T | -1.61 |
| M185L/K189T | -0.26 | C124A/D148K | -0.61 | C124A/K189S | -1.64 |
| C124A/G269W | -0.27 | A123S/C124A | -0.62 | A123S/C124G | -1.69 |
| M185L/G269V | -0.29 | C124A/K189T | -0.62 | C124G | -1.69 |
| M185L/F273W | -0.29 | C124G/K189L | -0.62 | M185L/K189A | -1.76 |
| K189G/N264K | -0.29 | C124T/K189S | -0.63 | A143M/K189A | -1.87 |
| C124G/M174L | -0.29 | C124G/L225I | -0.63 | C124A/K189A | -1.97 |
| C124G/K189E | -0.29 | A143M | -0.63 | C124G/M185L | -2.12 |
| C124L/K189A | -0.30 | K189S/F273W | -0.64 | C124G/A143M | -2.42 |

| | | | | | |
|---|---|---|---|---|---|
| C124G/N264S | -0.30 | C124P/K189A | -0.64 | C124G/K189S | -2.43 |
| C124A/K189- | -0.30 | D148R/K189G | -0.66 | C124G/K189G | -2.53 |
| K189G/G269W | -0.31 | C124A/D148R | -0.66 | C124G/K189A | -3.00 |
| W156F/K189A | -0.31 | K189A/G269W | -0.66 | | |

* Mutants evaluated in the experiment are highlighted in red.

**Table S5**. Variants with redesigned enzyme scaffold from the MaxEnt model

| mutant | $E(\boldsymbol{S})$ | mutant | $E(\boldsymbol{S})$ | mutant | $E(\boldsymbol{S})$ |
|---|---|---|---|---|---|
| Wild-type | 0.00 | F33K/K113N | -0.21 | K113N/Y298A | -0.62 |
| N35S/A43S | 0.00 | N35S/A200D | -0.21 | I75A/T102S | -0.62 |
| I75A/K209P | 0.00 | E40G/E195T | -0.21 | T102S/E195T | -0.62 |
| I67V/R305K | 0.00 | L250I/Y298A | -0.21 | F33K/E195S | -0.62 |
| L107T/Y298A | 0.00 | Y298A/R305E | -0.21 | V64I/T102S | -0.62 |
| A246R/Y298A | 0.00 | A246T | -0.21 | S130C/V212S | -0.63 |
| E195T/A200D | 0.00 | I74L/L107A | -0.21 | K113N/V212S | -0.63 |
| L107E/Y298A | -0.01 | E106D/Y298A | -0.21 | E195T/A246E | -0.63 |
| T102S | -0.01 | E40G/I75A | -0.22 | F33K/L107A | -0.64 |
| M27I/L107K | -0.01 | L107K/A246T | -0.22 | L107K/E195T | -0.64 |
| E44K/V212S | -0.01 | M27V/I67V | -0.22 | A43K/V212S | -0.64 |
| F116I/E195T | -0.01 | Y131N/Y298A | -0.22 | A139G/Y298A | -0.65 |
| M27I/I74L | -0.01 | I74L/R305K | -0.22 | L107Q/Y298A | -0.65 |
| L107Q/E195T | -0.01 | H62N | -0.22 | F33K/T102S | -0.66 |
| F33K/V212A | -0.01 | V212S/A246N | -0.22 | M27I/N35S | -0.66 |
| Y298A/R305- | -0.01 | T102S/A246E | -0.22 | I74L/E195T | -0.66 |
| V212S/R305- | -0.02 | A200E/Y298A | -0.22 | F116I/Y298A | -0.67 |
| V64I/V212A | -0.02 | E44G/Y298A | -0.23 | I34M/A246T | -0.67 |
| P69G/Y298A | -0.02 | H62N/T102S | -0.23 | V64I/R305K | -0.67 |
| F33K/A139G | -0.02 | A246T/R305K | -0.23 | I34M/H62N | -0.68 |
| L107A/K136R | -0.02 | V63I/V212S | -0.24 | I75A/L107A | -0.69 |
| E195T/A246K | -0.03 | F116L/Y298A | -0.24 | I34M/A246E | -0.70 |
| I74L/T102S | -0.03 | L107K/A246E | -0.24 | I74L/I75A | -0.70 |
| V212S/V303Y | -0.03 | Y131N/V212S | -0.24 | N35S | -0.71 |
| E195S | -0.04 | A46T/Y298A | -0.24 | V64I/E195S | -0.71 |
| I74L/L107K | -0.04 | V212S/L250I | -0.24 | L107Q/V212S | -0.72 |
| L107A | -0.04 | F33K/K136R | -0.25 | N35S/L107A | -0.73 |
| K136R/A246E | -0.04 | F33T/V212S | -0.25 | L107S/Y298A | -0.74 |
| A43Q/V212S | -0.04 | I34M/I74L | -0.25 | E195T/A246T | -0.74 |
| V70L/V212S | -0.04 | M27V | -0.25 | E195T/R305K | -0.74 |
| F33K/F116I | -0.04 | F33E/E195T | -0.25 | E40G/Y298A | -0.75 |
| A46P/V212S | -0.04 | F33E/V64I | -0.26 | N35S/T102S | -0.76 |
| V212S/E294A | -0.05 | T102S/A246T | -0.26 | N35S/R305K | -0.76 |
| L107K | -0.05 | V64I/A246K | -0.26 | V64I/I74L | -0.77 |

| | | | | | |
|---|---|---|---|---|---|
| T102S/L107A | -0.05 | I34M/R305K | -0.26 | F33E/Y298A | -0.77 |
| L107A/R305K | -0.05 | E44G/V212S | -0.26 | A246Q/Y298A | -0.78 |
| L107S/E195T | -0.05 | F33T/Y298A | -0.27 | V64I/L107K | -0.78 |
| P69G/V212S | -0.05 | N35S/F116I | -0.27 | F116I/V212S | -0.78 |
| E195T/R305S | -0.05 | I34M/I67V | -0.27 | N35S/E195S | -0.78 |
| I74L/K136R | -0.05 | M27L/V212S | -0.27 | I75A/A246T | -0.79 |
| I74L | -0.05 | K136R/E195T | -0.27 | F33K/A246T | -0.80 |
| I67V/I74L | -0.05 | I75A/A291S | -0.28 | H62N/I75A | -0.80 |
| N45D/V212S | -0.05 | E40G/V64I | -0.28 | F33K/R305K | -0.80 |
| M27I/A246T | -0.06 | H62N/L107A | -0.28 | F33K/L107K | -0.80 |
| M27V/A291S | -0.06 | E195S/A246E | -0.28 | I75A/A246E | -0.80 |
| V212T/Y298A | -0.06 | V64I/V303L | -0.28 | N35S/I74L | -0.81 |
| T102S/E195S | -0.06 | N35S/L107S | -0.29 | M27V/F33K | -0.81 |
| Y58F/V212S | -0.06 | I75A/L95V | -0.29 | H62N/E195T | -0.81 |
| I34M/E40G | -0.06 | V64I/A246Q | -0.29 | A246K/Y298A | -0.82 |
| K113N/E195T | -0.06 | I34M/L107A | -0.29 | M27V/I75A | -0.83 |
| G90I/Y298A | -0.07 | V212S/R305E | -0.29 | V212S/A246Q | -0.83 |
| N35S/S130C | -0.07 | F33K/V303L | -0.29 | F33K/A246E | -0.83 |
| I34M/K136R | -0.07 | V64I/A291S | -0.29 | V64I/L107A | -0.83 |
| L107K/R305K | -0.07 | E195D/V212S | -0.30 | N35S/L107K | -0.83 |
| Y129W/Y298A | -0.07 | E195T/A291S | -0.31 | L107S/V212S | -0.84 |
| V64I/L107Q | -0.07 | A246S/Y298A | -0.31 | K209P/V212S | -0.85 |
| N35S/L107Q | -0.08 | M27V/R305K | -0.31 | V64I/A246E | -0.85 |
| A43K/V64I | -0.08 | M27V/A246E | -0.31 | E40G/V212S | -0.86 |
| N35S/A43K | -0.08 | F33K/E40G | -0.31 | M27V/I34M | -0.86 |
| Y298A/S301D | -0.08 | L95V/E195T | -0.31 | I34M/E195T | -0.86 |
| I67V/L107K | -0.08 | M27V/E195S | -0.32 | M27V/E195T | -0.88 |
| I75A/F116I | -0.08 | E195S/A246T | -0.32 | A291S/Y298A | -0.89 |
| S87L/Y298A | -0.09 | V64I/L107S | -0.32 | V212S/V303L | -0.90 |
| F33K/R305S | -0.09 | A46T/V212S | -0.32 | V212S/A291S | -0.90 |
| I67V/A246T | -0.09 | V64I/L95V | -0.33 | L95V/V212S | -0.90 |
| R305K | -0.09 | E106D/V212S | -0.33 | N35S/A246T | -0.91 |
| L107E/V212S | -0.09 | M27I/H62N | -0.34 | V212S/A246K | -0.91 |
| N35S/A139G | -0.10 | F33K/L95V | -0.34 | V64I/A246T | -0.92 |
| M27V/L95V | -0.10 | M27V/L107K | -0.34 | F33K/H62N | -0.93 |
| H62N/K136R | -0.10 | E40D/V212S | -0.34 | F33E/V212S | -0.93 |

| | | | | | |
|---|---|---|---|---|---|
| T102D/Y298A | -0.10 | I75A/K136R | -0.35 | L95V/Y298A | -0.93 |
| I67V/L107A | -0.10 | F116L/V212S | -0.35 | N35S/A246E | -0.94 |
| I34M/A291S | -0.11 | N35S/A246Q | -0.35 | K136R/Y298A | -0.94 |
| V64I/R305S | -0.11 | M27V/T102S | -0.35 | M27V/V64I | -0.95 |
| V212S/A246R | -0.11 | I34M/E195S | -0.36 | H62N/V64I | -0.96 |
| V64I/K113N | -0.11 | H62N/L107K | -0.36 | N35S/H62N | -0.98 |
| T102S/L107K | -0.11 | N35S/V303L | -0.36 | K136R/V212S | -1.00 |
| T102S/R305K | -0.12 | I34M | -0.36 | F33K/I34M | -1.01 |
| N35S/V212D | -0.12 | N35S/A246K | -0.36 | I34M/I75A | -1.02 |
| M27I/R305K | -0.12 | H62N/A246T | -0.37 | I34M/V64I | -1.02 |
| N45D/Y298A | -0.12 | F33E/N35S | -0.37 | I75A/E195T | -1.04 |
| V64I/K209P | -0.12 | E195T/V303L | -0.37 | I34M/N35S | -1.04 |
| I34M/L95V | -0.12 | I34M/L107K | -0.37 | I67V/V212S | -1.09 |
| Y129W/V212S | -0.13 | N35S/E40G | -0.38 | M27V/N35S | -1.09 |
| I75A/V303L | -0.13 | V212S/A246S | -0.38 | I67V/Y298A | -1.11 |
| V212S/S301D | -0.13 | N35S/L95V | -0.38 | F33K/I75A | -1.13 |
| M27I/L107A | -0.13 | M27V/H62N | -0.39 | Y298A | -1.13 |
| N35S/R305S | -0.13 | H62N/E195S | -0.40 | M27I/Y298A | -1.16 |
| I137V/V212S | -0.13 | I34M/T102S | -0.40 | N35S/E195T | -1.16 |
| E195S/R305K | -0.13 | A43S/Y298A | -0.42 | E195S/Y298A | -1.17 |
| H62N/R305K | -0.13 | T102E/Y298A | -0.43 | M27I/V212S | -1.18 |
| F33K/A43K | -0.13 | A246D/Y298A | -0.44 | L107K/Y298A | -1.19 |
| I67V/A246E | -0.14 | I67V/I75A | -0.44 | T102S/Y298A | -1.19 |
| L107A/A246E | -0.14 | V212S/A246D | -0.44 | V212S | -1.21 |
| M27I/A246E | -0.14 | Y298A/R305S | -0.45 | Y298A/R305K | -1.22 |
| N35S/K209P | -0.14 | M27V/A246T | -0.46 | F33K/E195T | -1.23 |
| K136R/A246T | -0.14 | M27I/E195T | -0.46 | T102S/V212S | -1.25 |
| A246N/Y298A | -0.15 | V64I/K136R | -0.47 | F33K/V64I | -1.26 |
| F33K/A246K | -0.15 | A43K/Y298A | -0.48 | L107A/V212S | -1.28 |
| V64I/A200D | -0.15 | N35S/A291S | -0.48 | V64I/E195T | -1.28 |
| L107T/V212S | -0.15 | T102E/V212S | -0.48 | L107A/Y298A | -1.30 |
| N35S/V212A | -0.15 | N35S/K136R | -0.49 | A246E/Y298A | -1.31 |
| L107K/E195S | -0.16 | S130C/Y298A | -0.49 | I74L/Y298A | -1.31 |
| S87L/V212S | -0.16 | H62N/A246E | -0.49 | E195S/V212S | -1.32 |
| Y58F/Y298A | -0.16 | M27V/I74L | -0.50 | V212S/R305K | -1.33 |
| A246E | -0.16 | V212D/Y298A | -0.50 | I74L/V212S | -1.34 |

| | | | | | |
|---|---|---|---|---|---|
| V212G/Y298A | -0.16 | F33K/I74L | -0.50 | N35S/I75A | -1.34 |
| H62N/I74L | -0.16 | V64I/I67V | -0.50 | L107K/V212S | -1.35 |
| I75A/L107S | -0.16 | F33K/A291S | -0.51 | F33K/N35S | -1.35 |
| M27I/I34M | -0.16 | M27I/I75A | -0.52 | H62N/Y298A | -1.37 |
| I74L/A246T | -0.16 | A43S/V212S | -0.53 | N35S/V64I | -1.38 |
| M27L/Y298A | -0.17 | A139G/V212S | -0.53 | V212S/A246E | -1.40 |
| G90I/V212S | -0.17 | A200D/Y298A | -0.53 | V212S/A246T | -1.42 |
| M27V/K136R | -0.17 | V212A/Y298A | -0.54 | A246T/Y298A | -1.42 |
| I74L/E195S | -0.18 | M27I/V64I | -0.54 | M27V/Y298A | -1.43 |
| E195D/Y298A | -0.18 | L107A/E195T | -0.55 | M27V/V212S | -1.47 |
| E40D/Y298A | -0.18 | I75A | -0.55 | H62N/V212S | -1.48 |
| V64I/F116I | -0.18 | I75A/E195S | -0.55 | I34M/Y298A | -1.58 |
| F33K/A246Q | -0.18 | A200D/V212S | -0.55 | I34M/V212S | -1.60 |
| H62N/I67V | -0.19 | I67V/E195T | -0.56 | V64I/I75A | -1.66 |
| N35S/K113N | -0.19 | E195T | -0.56 | I75A/Y298A | -1.74 |
| I137V/Y298A | -0.19 | V212S/R305S | -0.58 | F33K/Y298A | -1.75 |
| A43D/V212S | -0.19 | K209P/Y298A | -0.58 | E195T/Y298A | -1.76 |
| T102D/V212S | -0.19 | Y298A/V303L | -0.58 | V64I/Y298A | -1.78 |
| I75A/A246K | -0.19 | I75A/L107K | -0.59 | E195T/V212S | -1.80 |
| L107A/A246T | -0.19 | I75A/R305K | -0.59 | F33K/V212S | -1.80 |
| A246E/R305K | -0.19 | M27V/L107A | -0.60 | I75A/V212S | -1.81 |
| F33K/L107S | -0.20 | V64I | -0.62 | N35S/Y298A | -1.86 |
| A200E/V212S | -0.20 | M27I/F33K | -0.62 | V64I/V212S | -1.91 |
| F33E/I75A | -0.20 | F33K/I67V | -0.62 | N35S/V212S | -1.97 |
| L107A/E195S | -0.20 | N35S/I67V | -0.62 | V212S/Y298A | -2.34 |
| I75A/A246Q | -0.20 | F33K | -0.62 | | |

* Mutants evaluated in the experiment are highlighted in red.

**Table S6**. Relative activity of RLuc variants tested in the experiment based on five measurements

| mutant | relative activity |
|---|---|
| Wild-type | 1.000 ± 0.066 |
| C124V | 0.006 ± 0.001 |
| M185L | 2.011 ± 0.140 |
| A143M | 1.020 ± 0.037 |
| K189S | 0.290 ± 0.015 |
| C124A | 1.631 ± 0.067 |
| K189G | 0.219 ± 0.006 |
| K189A | 0.293 ± 0.036 |
| C124G | 1.717 ± 0.069 |
| I34M | 0.604 ± 0.255 |
| I75A | 0.398 ± 0.079 |
| F33K | 0.648 ± 0.079 |
| V64I | 0.911 ± 0.076 |
| N35S | 0.329 ± 0.009 |
| V212S | 0.691 ± 0.197 |

**Table S7**. Stability of RLuc variants tested in the experiment based on three measurements

| mutant | $T_m$(°C) |
|---|---|
| Wild-type | 37.8 ± 0.2 |
| C124V | 39.3 ± 0.0 |
| M185L | 36.8 ± 0.0 |
| A143M | 40.2 ± 0.1 |
| K189S | 37.8 ± 0.0 |
| C124A | 40.4 ± 0.0 |
| K189G | 39.5 ± 0.0 |
| K189A | 39.2 ± 0.0 |
| C124G | 36.5 ± 0.0 |
| I34M | 35.6 ± 0.0 |
| I75A | 43.8 ± 0.1 |
| F33K | 40.9 ± 0.1 |
| V64I | 36.3 ± 0.2 |
| N35S | 32.8 ± 0.5 |
| V212S | 40.1 ± 0.0 |
